# Evolutionary rewiring of host metabolism and interferon signalling by SARS-CoV-2 variants

**DOI:** 10.1101/2025.11.15.688605

**Authors:** Riaz-Ali Somji, Jonathan C Brown, Shahid Rowles-Khalid, Tukur Abdullahi, Dongsheng Luo, Natalie Barthel, Michael Mülleder, Markus Ralser, Wendy S Barclay, Charalampos Rallis, Efstathios S Giotis

## Abstract

SARS-CoV-2 variants differ in transmissibility and immune evasion, but their effects on host-cell metabolism and signalling remain less defined. Using integrated transcriptomic, phosphoproteomic, and amino acid profiling in primary nasal epithelial cells, we compared early and late host responses to pre-Omicron variants (Alpha, Beta), Delta, and Omicron subvariants (BA.1, BA.5). Pre-Omicron strains broadly suppressed antiviral interferon-stimulated gene expression and reprogrammed metabolism by reducing mitochondrial oxidative phosphorylation and β-oxidation. Delta infection was associated with extensive transcriptional and metabolic remodelling, characterised by activation of stress- and growth-related kinases and selective retention of biosynthetic amino acids, consistent with a host response to stress and viral modulation of interferon-associated signalling. In contrast, Omicron infection elicited a more restrained response dominated by cytokine and survival pathways, with limited metabolic activation and interferon suppression. Together, these findings suggest SARS-CoV-2 has progressively evolved toward a strategy that maintains efficient upper-airway replication while minimising epithelial stress and inflammation.

## Introduction

SARS-CoV-2, the causative agent of COVID-19, has evolved into multiple variants with distinct transmissibility, immune evasion capacity, and associated disease severity. The successive waves of Variants of Concern (VOCs), including Alpha (PANGO lineage B.1.1.7), Beta (B.1.351), Delta (B.1.617.2/AY sublineages), and the many Omicron sublineages (B.1.1.529/BA sublineages, such as BA.1, BA.2, and BA.5), exhibit distinct phenotypic characteristics and transmission advantages over preceding variants. These advantages are largely driven by genetic changes in the viral genome, particularly in the spike protein, which enhanced receptor binding affinity and immune escape (Carabelli *et al*., 2023; Harvey *et al*., 2021; Liu and Rocklov, 2021). Additionally, VOCs exhibit varying capacities to suppress innate immune responses, a key determinant of early viral control and disease progression (Carabelli *et al*., 2023; Harvey *et al*., 2021; Laine *et al*., 2022; Reuschl *et al*., 2024).

The nasal epithelium serves as the primary entry site for SARS-CoV-2 infection and plays a critical role in shaping early host responses (Shi *et al*., 2024; Sungnak *et al*., 2020). Nasal epithelial cells (NECs) serve as both the primary site of viral entry and replication and as an essential frontline of antiviral immunity, producing interferons (IFNs), a family of cytokines that coordinate antiviral defence (Desmyter *et al*., 1968; Sposito *et al*., 2021). Type I (e.g., IFN-α and IFN-β) and type III (e.g., IFN-λ) interferons bind to their respective receptors on infected and neighbouring cells, triggering the Janus kinase/signal transducer and activator of transcription (JAK-STAT) pathway. Type II interferon (IFN-γ), primarily derived from immune effector cells, complements these pathways by amplifying antiviral and immunomodulatory responses once immune infiltration occurs. This leads to the induction of hundreds of interferon-stimulated genes (ISGs), which act collectively to inhibit viral replication, degrade viral RNA, and enhance antigen presentation (Park and Iwasaki, 2020; Schneider *et al*., 2014).

SARS-CoV-2 has evolved multiple strategies to suppress these IFN responses, thereby evading early immune detection and promoting viral persistence (Carabelli *et al*., 2023; V’Kovski *et al*., 2021; Yuen *et al*., 2020). The Alpha and Beta strains inhibit IFN production more effectively than earlier reference strains (Gori Savellini *et al*., 2023; Rajah *et al*., 2021). Delta, in contrast, displays enhanced replication efficiency and greater host immune antagonism, leading to higher viral loads and increased disease severity (Liu and Rocklov, 2021; Rajah *et al*., 2021). In comparison, several studies suggest that Omicron subvariants exhibit a reduced interaction with the interferon response, although some evidence indicates a delayed yet stronger induction of ISGs (Gori Savellini *et al*., 2023; Laine *et al*., 2022; Shi *et al*., 2024). Collectively, these findings suggest that each variant has evolved unique strategies to modulate IFN signalling and evade early antiviral defenses.

Among the emerging variants, Omicron variants remain globally dominant, as they rapidly displaced previous VOCs, through successive waves of transmission. Unlike earlier SARS-CoV-2 variants, Omicrons show an increased preference for the upper airways and a reduced transition to the lower respiratory tract (Shi *et al*., 2024). Studies suggest that mutations in the Omicron spike protein enhance viral entry into nasal epithelial cells, enabling evasion of both constitutive and interferon-induced antiviral factors that normally limit SARS-CoV-2 entry following attachment (Gori Savellini *et al*., 2023; Reuschl *et al*., 2024; Shi *et al*., 2024). In contrast to previous variants that predominantly rely on serine transmembrane proteases for host cell entry, Omicron strains preferentially use cysteine proteases cathepsin B (CatB) and cathepsin L (CatL) to catalyse endosomal membrane fusion, possibly providing an advantage in nasal epithelial infection and potentially contributing to their increased respiratory transmissibility (Giotis *et al*., 2022; Leach *et al*., 2021; Shi *et al*., 2024). These adaptations in entry mechanisms and tissue tropism raise important questions about how variant-specific traits impact downstream host cellular responses.

While extensive research on VOCs has focused on spike protein mutations and their role in enhancing viral entry and immune evasion, much less is known about how these variants differentially reprogram host cell metabolism and signalling to optimise replication and immune escape. Viruses such as Influenza A, Dengue, HBV, HIV and others typically hijack host metabolic networks to create an environment conducive to replication, altering energy production, amino acid biosynthesis, and lipid metabolism to support viral genome replication, protein synthesis, and virion assembly (Allen *et al*., 2022; Chan *et al*., 2009; Liu *et al*., 2020; Long *et al*., 2016; Polcicova *et al*., 2020; Ritter *et al*., 2010; Singh *et al*., 2020; Thaker *et al*., 2019). Current understanding of SARS-CoV-2 metabolic reprogramming largely derives from studies on early variants, which have shown that the virus targets key pathways such as glycolysis, the tricarboxylic acid (TCA) cycle, and the pentose phosphate pathway (PPP) to meet biosynthetic and redox demands (Codo *et al*., 2020; Guarnieri *et al*., 2024; Mullen *et al*., 2021). A central hallmark of this reprogramming leading to the accumulation of mitochondrial reactive oxygen species (mROS) and consequent oxidative stress is the inhibition of mitochondrial oxidative phosphorylation (OXPHOS) (Guarnieri *et al*., 2024). Elevated mROS stabilises hypoxia-inducible factor 1-alpha (HIF-1α), a key regulator that redirects cellular metabolism toward glycolysis and the PPP, enhancing the synthesis and mobilisation of nucleotides, lipids, and amino acids, which are necessary for viral replication (Guarnieri *et al*., 2024; Mullen *et al*., 2021). The virus also activates the mechanistic target of rapamycin complex 1 (mTORC1) and integrated stress response (ISR) pathways, which in-turn promote nutrient uptake, glycolysis, and innate immune signalling, while suppressing mitochondrial biogenesis and cytosolic protein synthesis (Guarnieri *et al*., 2024; Li *et al*., 2021; Mullen *et al*., 2021; Zambalde *et al*., 2022). In parallel, the virus disrupts cellular stress responses and modulates key signalling pathways, such as the phosphoinositide 3-kinase/protein kinase B (PI3K/AKT) and mitogen-activated protein kinase (MAPK) pathways, to promote cell survival (Bojkova *et al*., 2020; Li *et al*., 2021; V’Kovski *et al*., 2021). Despite these insights from early variants, how such metabolic and signalling circuits have been refined or rewired in later variants, particularly Delta and Omicron, remains poorly understood.

In this study, we focused on six epidemiologically and biologically representative SARS-CoV-2 lineages, which capture the principal stages of viral adaptation in replication dynamics, immune evasion, and host metabolic engagement during the pandemic. Using an integrated multi-omics approach in primary human nasal epithelial cells cultured at the air-liquid interface, we compared their transcriptional, signalling, and metabolic responses at early and late stages of infection to define lineage-specific strategies of host manipulation and their implications for viral fitness and adaptation.

## Results

Understanding how SARS-CoV-2 variants remodel host cell metabolism and signalling is critical to deciphering their evolutionary adaptations in transmissibility and immune evasion. To investigate these processes, primary nasal epithelial cells (NECs) from a pool of human donors were cultured at the air-liquid interface and infected with three classes of SARS-CoV-2 variants: early strains (IC19 (a lineage B.1 SARS-CoV-2 reference strain), the pre-Omicron: Alpha, Beta), Delta, and Omicron subvariants (BA.1, and BA.5). NECs were derived from three commercially available donor pools (MucilAir™, Epithelix), each comprising epithelial cells from 10-14 independent human donors. This pooled design ensured physiological relevance and helped minimise donor-specific variability across infection conditions. Viral replication kinetics were measured over 72 hours. Transcriptomic analysis (RNA-seq) was conducted at 24 and 72 hpi, and phosphoproteomic and amino acid profiling were performed at 24 hpi to characterise early post-translational and metabolic responses (Figure 1A).

**Figure 1.**
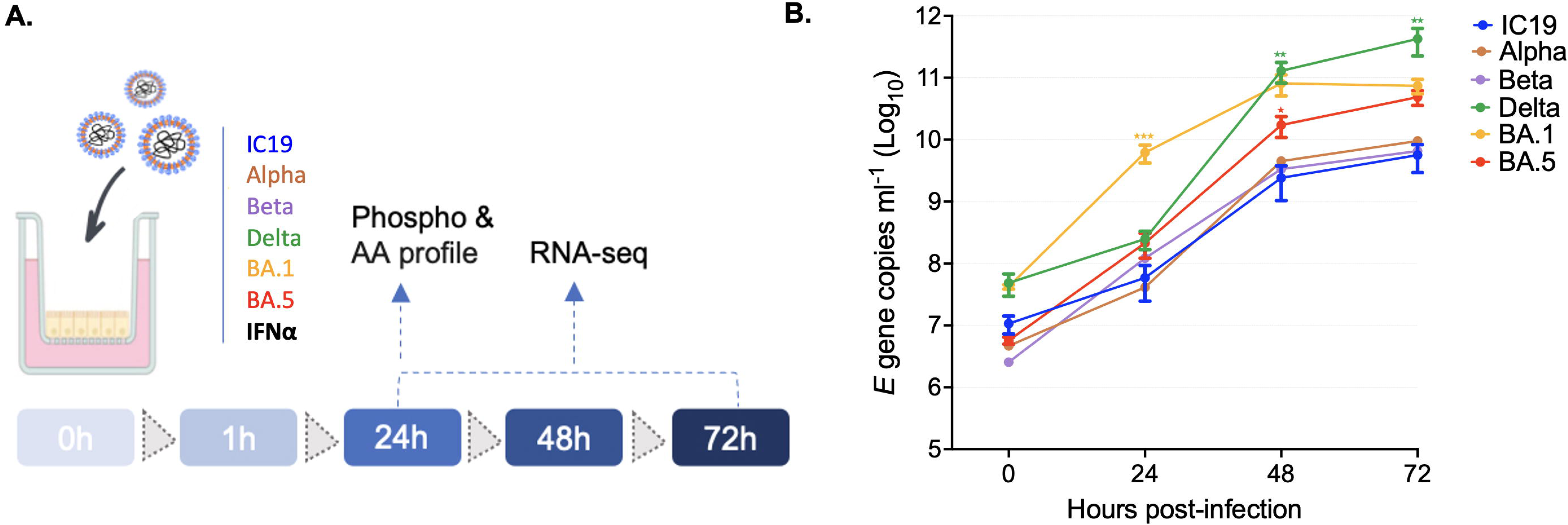
Omicron and Delta variants exhibit enhanced replication compared to pre-Omicron strains. **(A) Schematic representation of the experimental setup.** Primary human nasal epithelial cells were cultured at the air-liquid interface and infected with six SARS-CoV-2 variants: IC19 (reference strain), Alpha, Beta, Delta, Omicron BA.1, and Omicron BA.5. All infections were performed at a multiplicity of infection (MOI) of 0.01. Parallel wells treated with recombinant IFN-α served as a transcriptional benchmark. Cells were harvested at 24, 48, and 72 hpi for downstream analyses, including phosphoproteomic profiling, amino acid (AA) profiling, and RNA sequencing (RNA-seq). The timeline indicates the specific time points at which each molecular analysis was conducted. **(B) Growth kinetics of SARS-CoV-2 variants.** Viral replication over a 72-hour period was quantified by RT-qPCR targeting the *E* gene. Viral RNA levels are expressed as *E* gene copies per millilitre on a log10 scale. Measurements were taken at 0, 24, 48, and 72 hpi. Variants Delta and BA.1 demonstrated significantly elevated viral loads compared to the reference strain IC19 at multiple time points. Statistical significance was evaluated using one-way ANOVA with Tukey’s multiple comparisons test. Asterisks indicate significance levels: *p < 0.05, **p < 0.01, ***p < 0.001. Data are shown as mean ± standard deviation (SD) from three independent experiments.

### Omicron and Delta variants exhibit increased replication compared to pre-Omicron strains

To compare the replication dynamics of SARS-CoV-2 variants, NECs were infected with the variants at a multiplicity of infection (MOI) of 0.01, and viral genome copies in the culture supernatants were quantified over time using *E* gene RT-qPCR (Figure 1B).

At 24 hpi, BA.1 exhibited the highest early replication, with significantly elevated viral genome copies compared to the pre-Omicron reference strain IC19 (*p* < 0.0001), indicating an early replication advantage. By 48 and 72 hpi, Delta showed the highest viral loads, significantly exceeding IC19 at both time points (*p* = 0.0014 and 0.0029, respectively). BA.5 and Beta also demonstrated increased replication relative to IC19, with significantly higher viral titres at 48 hpi (*p* = 0.0165 and 0.0173) and 72 hpi (*p* = 0.0261 and 0.0288). These findings demonstrate group-specific replication dynamics, with Delta and Omicron subvariants exhibiting increased replication efficiency in nasal epithelial cells compared to pre-Omicron strains.

### Delta triggers the most extensive transcriptional remodelling late in infection

To characterise host transcriptional responses to SARS-CoV-2 variants, we performed RNA-seq on NECs at 24 and 72 hpi (Figure 2A-C). Differentially expressed genes (DEGs) were defined by a log₂ fold change ≥ |1| and FDR ≤ 0.05 (Table S1). Key RNA-seq findings were validated by qPCR for representative immune-signalling and metabolic genes (Supplementary Figures S1–S2).

**Figure 2.**
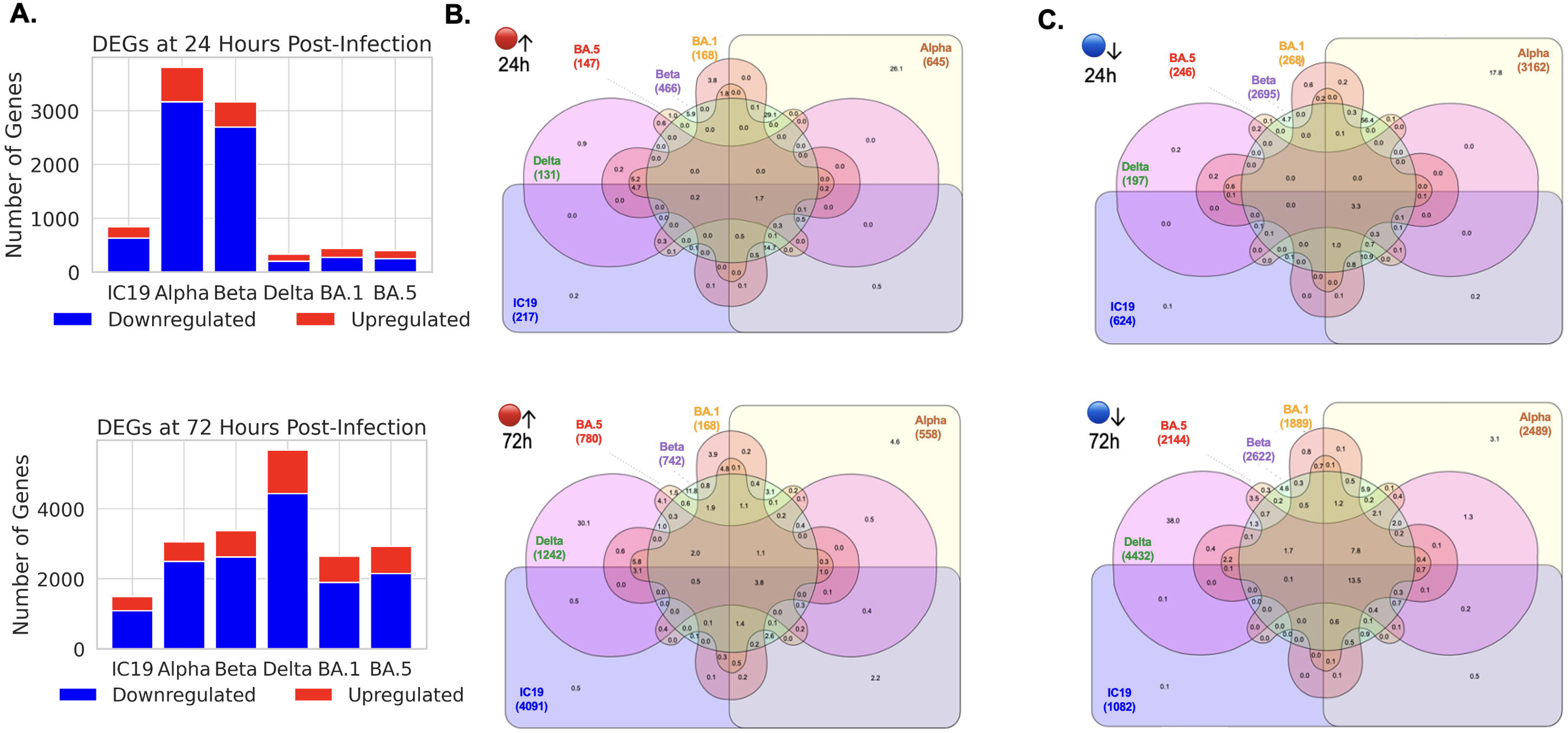
Transcriptomic profiling of host responses to SARS-CoV-2 variant infection. (A) Number of significantly up-regulated (red) and down-regulated (blue) differentially expressed genes (DEGs) at 24 and 72 hpi with indicated SARS-CoV-2 variants, based on RNA-seq analysis (log2 fold-change ≥1; FDR <0.05). (B) Venn diagrams showing shared and unique up-regulated DEGs among variants at 24 hpi (top) and 72 hpi (bottom). (C) Venn diagrams showing shared and unique down-regulated DEGs among variants at 24 hpi (top) and 72 hpi (bottom). Percentages represent overlap relative to the total DEGs per variant.

At 24 hpi, pre-Omicron variants (IC19, Alpha, and Beta) elicited the most substantial early responses, with Alpha alone triggering over 3,800 DEGs (645 up-regulated, 3,162 down-regulated). In contrast, Delta and the Omicron subvariants (BA.1 and BA.5) triggered markedly fewer changes, suggesting attenuated or delayed early sensing (Figure 2B).

By 72 hpi, Delta drove the most extensive transcriptional reprogramming across all variants (1,242 up-regulated, 4,432 down-regulated genes), while BA.1 and BA.5 induced intermediate responses. Although some overlap in down-regulated genes was observed between Delta and Omicron, the majority of DEGs remained variant-specific (Figure 2C).

Together, these data reveal distinct temporal transcriptional patterns across SARS-CoV-2 variants, with Delta driving the broadest late-stage reprogramming and highlighting the evolving diversity of host–virus interactions.

### Reduced suppression of host interferon responses following infection with Omicron subvariants

We next examined how SARS-CoV-2 variants modulate innate antiviral signalling by measuring interferon-stimulated gene (ISG) expression in primary nasal epithelial cells at 24 and 72 hpi (Figure 3A–D). ISGs represent core effectors of the type I interferon pathway and provide a transcriptional readout of antiviral activation (Park and Iwasaki, 2020) (Figure 3A). Cells treated with recombinant interferon alpha (IFN-α) were included as a reference to establish a benchmark for ISG induction.

**Figure 3.**
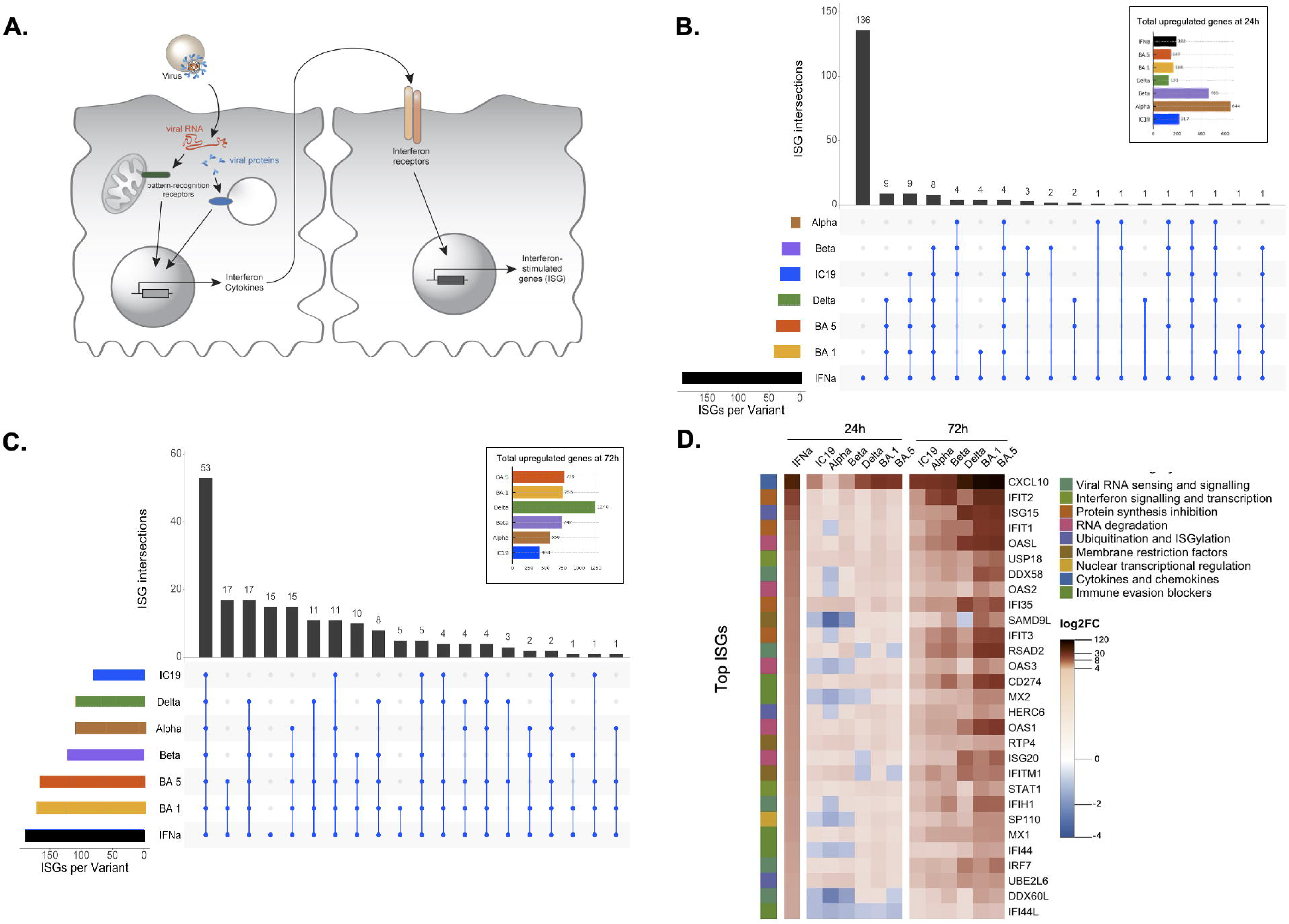
Late ISG induction by Omicron strains versus sustained attenuation by Delta. (**A**) **Schematic of interferon signalling and ISG activation.** (**B-C**) **Shared and unique ISG induction across variants at 24 (B) and 72 (C) hpi.** An UpSet plot visualizes the number of significantly up-regulated ISGs (FDR < 0.05) per SARS-CoV-2 variant and their overlaps. The bar chart at the top indicates the size of each intersection set, while the connected dots below identify which variants contribute to each group. The colored horizontal bars represent total ISG numbers detected per variant. The inset bar graphs show the total number of significantly upregulated genes detected for each variant at the corresponding time point, providing context for overall transcriptional activation. (**D**) **Heatmaps showing log2 fold-change of top ISGs induced by IFNα treatment and during infection at 24 and 72 hpi.** The leftmost panel shows the top ISGs up-regulated by IFNα treatment alone, identifying baseline interferon responsiveness of the cells. The middle and right panels show the expression of these ISGs in response to viral infection across all six SARS-CoV-2 variants at 24 and 72 hpi, respectively. Color intensity reflects log2 fold-change, with red indicating upregulation and blue indicating down-regulation or suppression.

At 24 hpi, ISG induction remained low across all variant-infected relative to mock samples (Figure 3B). Alpha and Beta showed the weakest early responses, whereas Delta and the Omicron subvariants BA.1 and BA.5 triggered modest but detectable up-regulation. By 72 hpi, ISG expression was broadly elevated, but with clear variant-specific differences (Figure 3C).

BA.1 and BA.5 induced the largest and most diverse ISG repertoires, similar in scale to those induced by IFN-α, while Alpha and Beta exhibited more restricted activation. In contrast, Delta consistently induced the fewest ISGs, even as infection advanced, pointing to sustained suppression of interferon responses.

A heatmap of the top IFN-α-responsive ISGs (Figure 3D) further illustrates these differences. BA.1 and BA.5 displayed relatively higher expression of canonical antiviral genes by 72 hpi such as *CXCL10*, *ISG15*, *IFIT1*, and *OAS1*, key effectors of RNA sensing, translational arrest, and immune recruitment (Schneider *et al*., 2014; Schoggins, 2019; Schoggins *et al*., 2011), compared with Delta. Compared with BA.1- and BA.5-infected cells, Delta-infected cells showed fewer IFN-α–responsive genes and reduced expression levels, suggesting a weaker activation of interferon signalling.

### Progressive tuning of host metabolism from early lineages to Delta and Omicron

Previous reports have shown that early SARS-CoV-2 lineages, including IC19, Alpha, and Beta, are associated with reduced mitochondrial activity and enhanced glycolytic flux, an established viral strategy to redirect cellular resources toward biosynthesis (Codo *et al*., 2020; Guarnieri *et al*., 2024; Mullen *et al*., 2021). Cytosolic pathways such as glycolysis and the pentose phosphate pathway (PPP) generate biosynthetic precursors, whereas mitochondrial metabolism, including the tricarboxylic-acid (TCA) cycle, fatty-acid oxidation (FAO), and oxidative phosphorylation (OXPHOS), supports ATP production, redox balance, and antiviral signalling.

To investigate how infection with different SARS-CoV-2 variants correlates with metabolic pathway changes in host cells, we performed gene set enrichment analysis (GSEA) of RNA-seq data from NECs infected with the variants at 24 and 72 hpi, relative to uninfected controls (Figure 4A-B). Normalised enrichment scores (NES) were used to quantify coordinated transcriptional changes across curated metabolic pathways, providing a comparative measure of pathway activation (positive NES) or suppression (negative NES) independent of gene-set size and variance (Figure 4; full data is listed in Table S2). For interpretation, pathways with an FDR < 0.05 were categorised by NES magnitude as follows: |NES| < 1.0, limited enrichment; 1.0 ≤ |NES| < 1.8, moderate enrichment; |NES| ≥ 1.8, strong enrichment.

**Figure 4.**
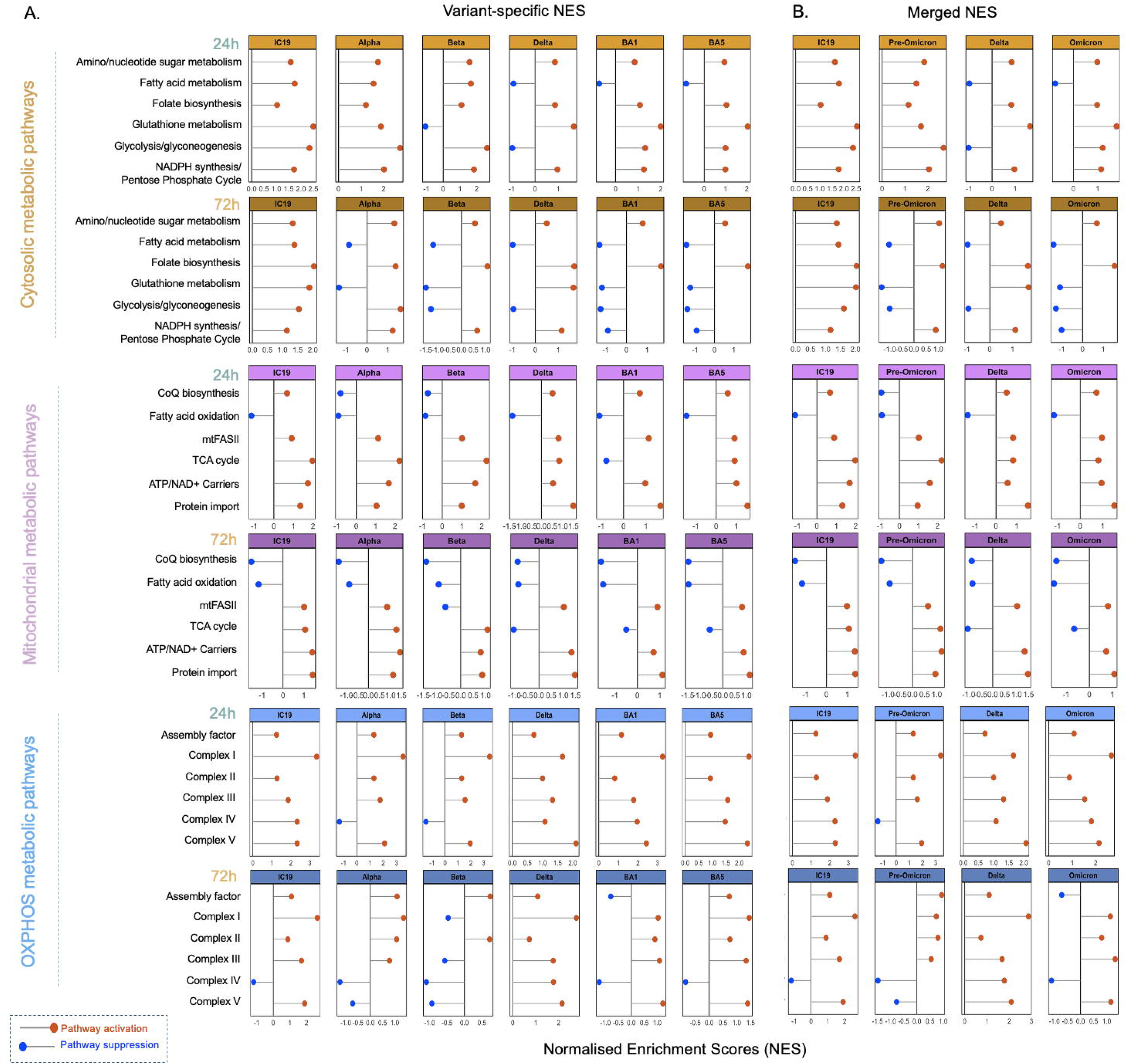
Normalised enrichment scores (NES) depict pathway-level transcriptional changes across SARS-CoV-2 variants and time points. (**A**) Variant-specific NES profiles for cytosolic, mitochondrial, and oxidative phosphorylation (OXPHOS) pathways at 24 and 72 hpi. Differential expression values (log₂ fold change) were ranked and analysed with the fgsea R package using curated KEGG and literature-based gene sets. (**B**) Merged analysis comparing Pre-Omicron and Omicron lineages, showing composite NES values summarizing transcriptional regulation across variants. Red indicates pathway activation, blue indicates suppression. NES, normalised enrichment score.

At 24 hpi, metabolic responses were limited in magnitude (Figure 4A). Cells infected with most variants showed near-neutral or mildly positive NES values in glycolysis and PPP, suggesting modest early cytosolic engagement. FAO gene sets were consistently negative across lineages, indicating an early block in β-oxidation. Mitochondrial carrier and import modules were close to neutral, while OXPHOS complexes displayed small but generally positive enrichment, apart from a modest negative enrichment in the Alpha/Beta Complex IV module. Thus, early infection was associated with preserved or moderately enhanced respiratory-gene expression rather than broad suppression.

By 72 hpi, clear lineage-specific profiles emerged (Figure 4B). Delta-infected cells exhibited the highest and most coordinated positive NES across OXPHOS complexes I–V and several mitochondrial modules, indicating strong respiratory activation accompanied by moderate enrichment of glycolytic and lipid-metabolism pathways. Cells infected with the reference strain IC19 retained moderate OXPHOS activation. Cells infected with pre-Omicron strains displayed overall weaker enrichment and more variable mitochondrial responses. In contrast, Omicron-infected cells showed near-neutral cytosolic NES, negative FAO and coenzyme Q biosynthesis scores, and modest positive enrichment of mitochondrial-carrier and OXPHOS modules, again with a negative Complex IV component. These findings suggest that Omicron infection elicits a host response that maintains basal mitochondrial function while avoiding extensive metabolic activation.

Grouped comparison (Figure 4B) summarises these three variant-specific trends. Cells infected with early lineages (IC19, Alpha, Beta) show moderate OXPHOS enrichment at 24 hpi that diminishes and becomes variable by 72 hpi. Delta-infected cells display the strongest and most coordinated activation across respiratory and associated metabolic pathways. Omicron-infected cells retain a selective, mitochondria-centred profile with positive overall OXPHOS and minimal cytosolic activation.

### Delta activates stress kinases, while Omicron reprograms cytokine and survival signalling

To characterise early host signalling responses, we next profiled phosphorylation events at 24 hpi in NECs infected with IC19, Delta, BA.1, and BA.5 using phospho-kinase arrays. These variants were selected to represent the most distinct host-response patterns observed in our dataset: IC19 as the ancestral reference, Delta as the lineage that showed the broadest transcriptional and metabolic engagement in infected cells, and BA.1/BA.5 as Omicron subvariants that, in our system, elicited comparatively restrained early responses but clearer cytokine-linked signalling at later stages. Phosphorylation levels were normalised to mock controls, and statistical significance was assessed using Welch’s *t*-tests with Benjamini–Hochberg correction (FDR < 0.05; Table S3).

Relative to mock, all infections were associated with phosphorylation changes in stress-, growth-, and cytokine-associated proteins, but the magnitude and pattern differed markedly between variants (Figure 5A–B). Compared with IC19-, Delta-infected cells displayed the most extensive activation of cellular stress and translational pathways, with significant increases in p53, PRAS40, and p70 kinase alongside higher phosphorylation of STAT3 and STAT6. This profile suggests enhanced engagement of stress-responsive and translational signalling pathways. Collectively, the pattern observed in Delta-infected cells is consistent with a stress-associated and pro-survival phosphorylation signature dominated by the p53–RSK–S6K and PRAS40 nodes.

**Figure 5.**
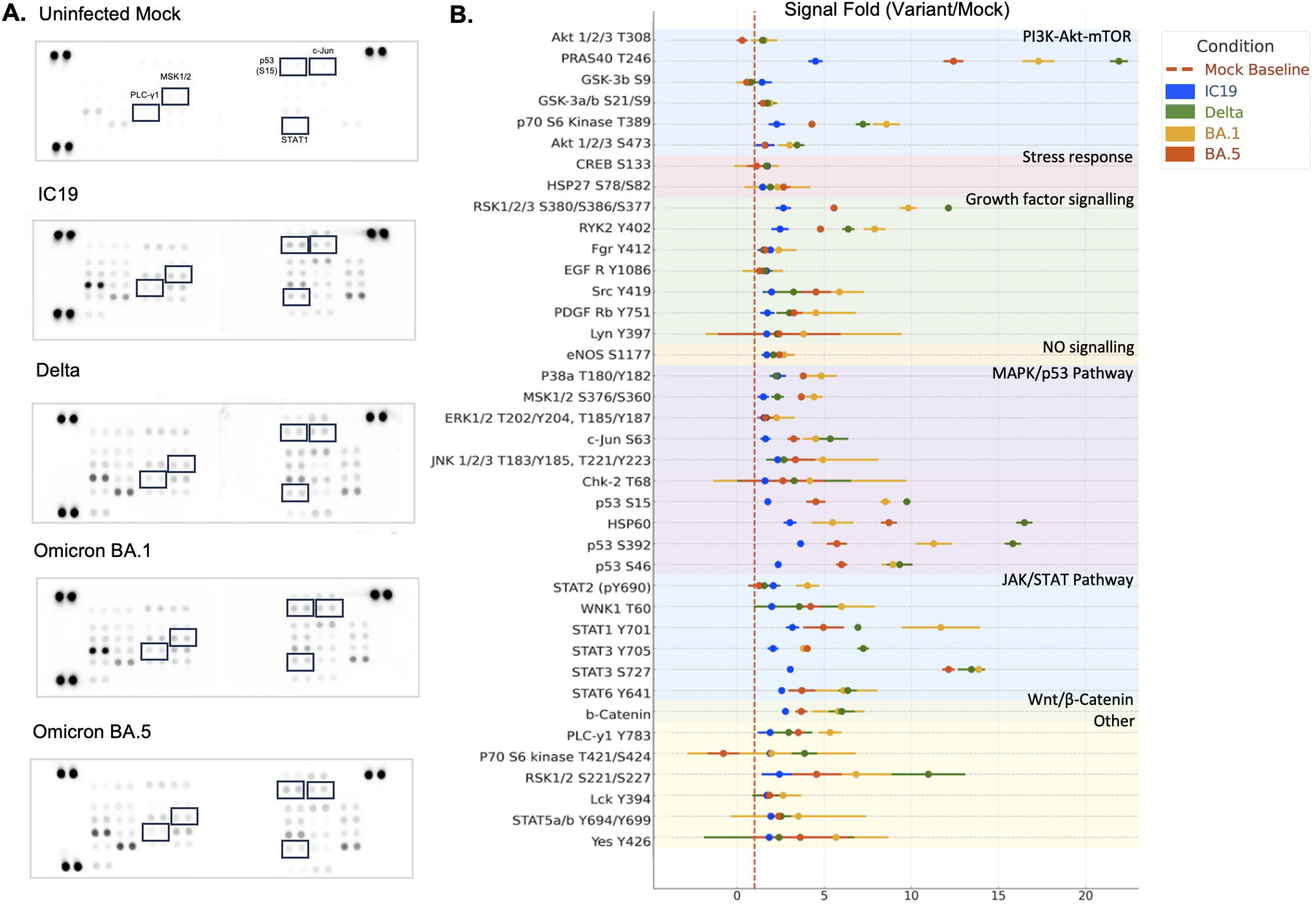
SARS-CoV-2 variants differentially regulate host signalling pathways through phosphorylation events. (**A**) **Representative phospho-protein arrays** showing phosphorylation patterns in uninfected mock controls and cells infected with IC19, Delta, Omicron BA.1, and Omicron BA.5 at 24 hpi. Key differentially phosphorylated proteins are highlighted. (**B**) **Quantitative analysis of phosphorylation fold changes (variant-infected vs. mock) across all array targets grouped by signalling pathway.** Dots represent mean values from at least two independent experiments, with vertical error bars showing the standard error of the mean (SEM). Colour coding denotes variant groups: IC19 (blue), Delta (orange), BA.1 (green), and BA.5 (red). Markers are ranked by biological pathway, with coloured background shading corresponding to biological grouping. The vertical red dashed line indicates the mock-infected baseline (fold change = 1).

By contrast, cells infected with Omicron subvariants exchibited phosphorylation patters enriched in immune- and cytokine-linked pathways. BA.1-infected cells showed prominent phosphorylation of p38 MAPK, PLC-γ1, RYK, and Src, alongside strong activation of JAK–STAT components including STAT1, STAT2, and STAT5. Phosphorylation fold changes for STAT1 (11.7-fold) and STAT2 (4.0-fold) were the highest among all variants, consistent with strong interferon- and cytokine-associated signalling. Cells infected with BA.5 showed a similar but overall attenuated profile compared with BA.1, retaining phosphorylation of the p53, STAT3 (S727), and Src-family kinases.

### Contrasting host amino acid regulation by Delta and Omicron variants

To complement the transcriptomic and phosphoproteomic analyses, intracellular amino acid levels were quantified at 24 hpi in nasal epithelial cells infected with IC19, Delta, BA.1 or BA.5 and compared with mock-infected controls (Figure 6; Table S4). Total intracellular amino acid abundance (Figure 6A) showed modest, non-significant reductions in IC19- and Delta-infected cells (*q* > 0.05), despite an approximate 10% decrease in IC19 relative to mock. In contrast, BA.1- and BA.5-infected cells exhibited significant reductions in total amino acid pools (*q* < 0.01 and *q* < 0.05, respectively). All amino acid measurements were expressed as nmol per 10⁶ viable, counted cells, and viability at 24 hpi was comparable across conditions, indicating that differences in abundance reflect infection-driven metabolic changes rather than variation in cell number.

**Figure 6.**
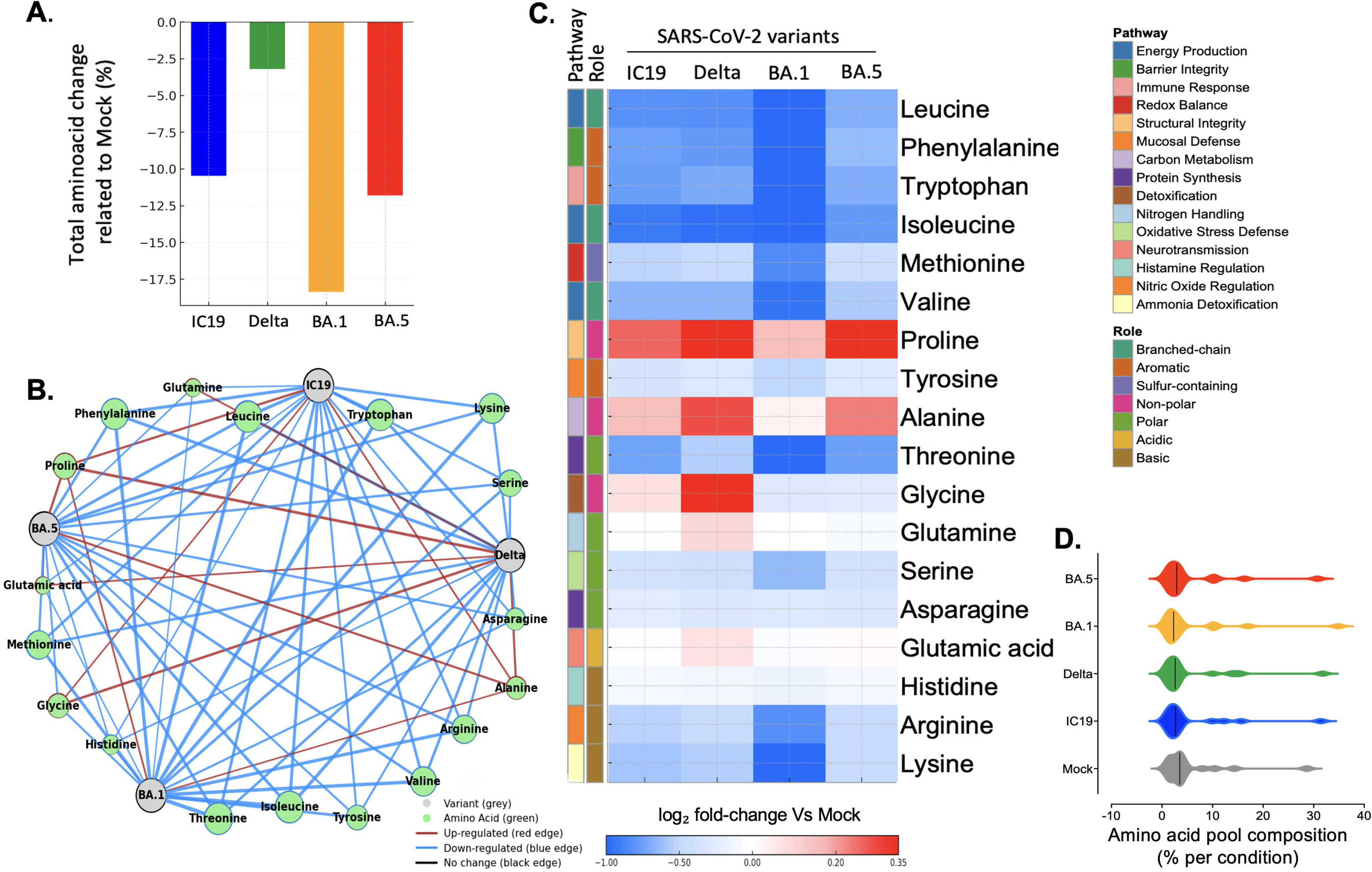
SARS-CoV-2 variants induce distinct amino acid reprogramming in nasal epithelial cells. (**A**) **Bar plot showing total amino acid abundance** (nmol/10⁶ cells) relative to mock-infected controls at 24 hpi. (**B**) **Variant–metabolite interaction network** showing amino acids (green nodes) connected to each SARS-CoV-2 variant (grey nodes). Edges represent directionality of regulation compared to mock: red edges indicate increased levels (upregulation), blue edges indicate decreased levels (down-regulation), and black edges indicate no change. (**C**) **Heatmap of log₂-transformed amino acid levels** (based on nmol/10⁶ cells) for each variant relative to mock-infected controls at 24 hpi. for each variant relative to mock-infected controls at 24 hpi. Amino acids are annotated by metabolic pathway (left colour bar) and biochemical role (right colour bar). Red shading indicates increased amino acid abundance, and blue shading indicates decreased abundance. **(D) Violin plots showing the % relative amino acid composition** after normalisation of each amino acid to the total amino acid content within the same condition. This representation highlights the internal proportional balance of amino acids rather than total abundance. The black line indicates the median value for each distribution.

The variant–amino acid interaction network (Figure 6B) visualises these differences by linking each variant to its most strongly altered amino acids. Edge width reflects the absolute log₂ fold change, and colour indicates direction (red for increases, blue for decreases relative to mock). BA.1 and BA.5 display numerous, broad negative edges, while IC19 and Delta show fewer, thinner connections, consistent with a more limited metabolic impact.

Individual amino acid profiles (Figure 6C; Table S4) further resolved these patterns. In BA.1-infected cells, intracellular amino acid levels showed the broadest and most significant reductions, including all branched-chain and aromatic amino acids (leucine, phenylalanine, tryptophan, isoleucine, valine), together with decreases in several other amino acids, including threonine, serine, arginine and lysine. BA.5-infected cells exhibited a similar but less pronounced profile, with significant decreases in many essential amino acids and moderate increases in alanine and proline. Delta-infected cells displayed a mixed pattern, characterised by significant increases in proline, alanine, glycine and glutamine, alongside reductions in selected essential amino acids such as isoleucine and phenylalanine. IC19-infected cells showed only small, variable changes, none of which reached significance after FDR correction. To examine whether infection altered the internal balance of amino acids rather than total abundance, amino acid levels were normalised to the total amino acid content within each condition (Figure 6D; Table S4). The relative amino acid composition appeared consistent across variants, suggesting that infection reduces overall amino acid availability without disturbing the proportional balance within the intracellular pool.

Together, these data show that amino acid abundances in variant-infected cells differ from those in mock-infected cells in a lineage-specific manner, with Delta-infected cells showing comparatively higher levels of several biosynthetically relevant amino acids, whereas cells infected with Omicron subvariants display broad amino acid depletion and limited compensatory increases across pathways. Amino acids are central to energy production, redox balance, protein synthesis, and immune signalling, and their intracellular abundance provides a sensitive readout of metabolic stress and viral resource use (Thaker *et al*., 2019).

## Discussion

SARS-CoV-2 variants have evolved distinct capacities to replicate, modulate host immunity, and influence disease severity. Although mechanisms of viral entry and antibody escape are well characterised, far less is understood about how major lineages remodel host cellular pathways once infection is established. To address this, we employed differentiated nasal epithelial cells (NECs) grown at air-liquid interface, a physiologically relevant model that recapitulates early epithelial infection in the absence of systemic immune influences. This system enables direct examination of variant-specific host responses at the primary site of replication, which may differ in magnitude and kinetics from those observed in bronchial or immune-cell containing models. Through an integrated multi-omics approach, we uncovered lineage-specific differences in the extent, timing, and coordination of transcriptional, signalling, and metabolic responses.

Early lineages (IC19, Alpha, Beta) broadly down-regulated immune and metabolic transcripts at 24 hpi, consistent with potent suppression of innate antiviral signalling. These results align with previous studies showing that early SARS-CoV-2 isolates suppress mRNA synthesis and translation through ORF6-, ORF8-, and ORF9b-mediated interference with STAT1/2 and nuclear import (Laine *et al*., 2022; Reuschl *et al*., 2024; V’Kovski *et al*., 2021; Yuen *et al*., 2020). By 72 hpi, infected nasal epithelial cells exhibited clear lineage-specific divergence. Cells infected with Delta exhibited extensive transcriptional reprogramming, including activation of stress-response, metabolic, and cytokine-related genes. In contrast, Omicron-infected cells displayed fewer differentially expressed genes, nevertheless, showed selective enrichment of interferon-stimulated and cytokine-responsive transcripts, consistent with a delayed yet measurable antiviral response (Gori Savellini *et al*., 2023; Reuschl *et al*., 2024; Shi *et al*., 2024). Consistent with previously reported effective antagonism of RIG-I/MDA5–JAK–STAT signalling (Laine *et al*., 2022; Sievers *et al*., 2024; Tandel *et al*., 2022) IC19, Alpha, Beta, and Delta displayed weak ISG induction at both 24 and 72 hpi. In contrast, Omicron elicited a broader ISG response by 72 hpi, approaching that observed following exogenous IFN-α treatment, suggesting only partial suppression of interferon-driven transcription (Reuschl *et al*., 2024; Shi *et al*., 2024).

Phosphoproteomic profiling further distinguished variant phenotypes. Omicron infections enhanced phosphorylation within JAK–STAT and MAPK modules (STAT1/2/5, p38, PLC-γ1, Src), while Delta preferentially activated stress- and growth-associated kinases (p53, PRAS40, RSK, p70 S6K), indicating engagement of PI3K–Akt–mTORC1 and the integrated stress response (Bouhaddou *et al*., 2020; Klann *et al*., 2020; Li *et al*., 2021). These patterns suggest that Omicron can replicate despite ongoing cytokine signalling, whereas Delta infection favours stress- and growth-linked pathways to sustain replication.

Our metabolic data reveal pronounced lineage-specific differences in host–virus interaction. Consistent with previous reports (Codo *et al*., 2020; Mullen *et al*., 2021), cells infected with pre-Omicron lineages suppressed fatty-acid oxidation and exhibited modest, heterogenous mitochondrial changes, indicating partial mitochondrial adaptation rather than broad inhibition. Conversely, Delta-infected cells displayed the strongest and most coordinated activation spanning oxidative-phosphorylation complexes I–V and several mitochondrial modules, accompanied by moderate enrichment of cytosolic pathways, including the tricarboxylic-acid cycle and lipid metabolism. These transcriptional shifts occurred despite broadly comparable viral loads across lineages at 24-72 hpi, suggesting that replication level alone does not account for the observed metabolic divergence. Delta-infected NECs retained or modestly increased one-carbon intermediates such as glycine, glutamine, and alanine, consistent with preserved anabolic and redox capacity. Targeted metabolomic analysis further revealed that total intracellular amino acid pools declined across infections relative to mock, but the extent varied by lineage: Delta-infected cells exhibited the smallest reduction, BA.5-infected cells an intermediate profile, and BA.1-infected cells the most pronounced depletion, particularly of essential and branched-chain amino acids such as leucine and valine. These findings indicate that Delta sustains a more biosynthetically active state, whereas Omicron subvariants replicate under more metabolically constrained conditions, suggesting a shift towards a low-anabolism infection strategy. When amino acid levels were normalised to total intracellular content, the overall proportional balance of amino acids in variant-infected cells was largely preserved, with only a small deviation detected for BA.1 compared with mock. This pattern suggests that SARS-CoV-2 lineages primarily modulate the metabolic tone of nasal epithelial cells by altering amino acid availability, rather than by extensively reshaping the internal composition of the amino acid pool.

Integration of transcriptomic, phosphoproteomic, and metabolic datasets, delineates three distinct host-virus interaction strategies. Early lineages dampened transcription, interferon signalling, and fatty-acid oxidation while maintaining only minimal mitochondrial activity. Delta, by contrast, exhibited coordinated activation of respiratory, cytosolic, and stress pathways, consistent with higher metabolic demand. Omicron retained efficient replication while preserving mitochondrial respiration and tolerating interferon-linked signalling, yet engaging anabolic metabolism only weakly. These lineage-specific programmes likely reflect not only viral genetic differences but also the interplay between replication dynamics and epithelial sensing thresholds in NECs. Overall, SARS-CoV-2 evolution appears to have progressively re-tuned the balance between replication, metabolism, and immune modulation. Later variants replicate effectively while imposing a smaller metabolic load and causing less epithelial stress, yet remain capable of replication in the presence of active cytokine and interferon responses. This restrained, energy-efficient infection profile mirrors clinical observations that Omicron replication is largely confined to the upper airways and associated with reduced epithelial injury and inflammation (Carabelli *et al*., 2023; Gori Savellini *et al*., 2023; Laine *et al*., 2022).

The use of pooled donor–derived NECs enhances physiological relevance but may mask inter-individual variation in antiviral and metabolic responses. Future studies sampling additional time points and integrating isotope-tracing could define metabolic flux through glycolysis and the TCA cycle, while targeted perturbation of pathways such as PI3K–Akt–mTORC1, p53–RSK–S6K, and p38 MAPK in ALI cultures could establish causal links between signalling nodes and replication outcomes. Extending the approach to lower-airway or immune–epithelial co-culture models would further connect cellular rewiring to tissue-level phenotypes and transmissibility.

In summary, SARS-CoV-2 variants employ distinct reprogramming strategies in nasal epithelium. Pre-Omicron strains strongly suppress antiviral and metabolic responses; Delta drives extensive activation of stress and energy pathways; Omicron maintains mitochondrial activity while limiting broader metabolic engagement. These contrasting strategies illustrate how viral evolution has refined replication efficiency relative to metabolic demand and immune modulation, providing mechanistic insight for host-targeted therapeutic approaches.

## Resource Availability Lead Contact

Further information and requests for resources and reagents should be directed to and will be fulfilled by the Lead Contact, Dr Efstathios Giotis.

## Materials availability

No new unique reagents or biological materials were generated in this study. SARS-CoV-2 variant strains were obtained from existing publicly available virus collections, as indicated in the Materials and Methods. All cell lines and virus isolates used are commercially or institutionally available.

## Data and code availability

The RNA sequencing data generated during this study have been deposited in the Gene Expression Omnibus (GEO) under accession number GSE271378. The phosphoproteomic and amino acid profiling datasets are available in the Supplemental information and can also be shared upon request from the Lead Contact. This study generated analysis scripts for figure production, available upon reasonable request from the Lead Contact. Scripts for data visualisation (e.g., R scripts for heatmaps, Python for statistical plots) are available from the Lead Contact upon reasonable request.

## Author contributions

E.S.G. conceptualised the study, supervised the project, and contributed to methodology, investigation, formal analysis, visualisation, and funding acquisition. R.A.S., J.B., T.A., N.B., D.L. and M.Mu. performed experimental work and contributed to methodology and investigation. S.R.K. carried out bioinformatic analyses, formal analysis, data curation, and visualisation. R.A.S. also contributed to data visualisation. M.Ra. conducted amino acid profiling and provided resources and supervision. W.S.B. and C.R. contributed to project supervision, resources, and funding acquisition. E.S.G., R.A.S., and S.R.K. wrote the original draft, and all authors reviewed and edited the manuscript.

## Acknowledgements

We are grateful to Juhi Kumar, Dorothee Reuss, Natalia Paulina Twardowska and Tom Peacock for their technical assistance and helpful input during manuscript preparation. This work was supported by the University of Essex COVID-19 Rapid and Agile Fund and the Faculty of Science and Health Research Innovation and Support Fund. EG and DL are funded by an MRC New Investigator Research Grant (MR/Z506242/1) and the Royal Society (RGS\R2\242527). C.R. acknowledges support and funding from the Biotechnology and Biological Sciences Research Council [Research grant numbers: BB/V006916/1, BB/V006916/2] and from the Medical Research Council [Grant number: MR/W001462/1]. Wendy Barclay is supported in part by the NIHR Imperial Biomedical Research Centre (BRC). Tukur Abdullahi is supported by the Nigerian Petroleum Technology Development Fund and the University of Essex.

## Supplemental information

**Supplementary Figures S1–S2.**

Contains validation of RNA-seq data by qPCR across selected immune signalling and metabolic genes. Includes:

- **Supplementary Figure S1:** Correlation plot of RNA-seq vs qPCR fold-change values for immune-signalling genes (*CXCL10, IFIT1, IFIT2, ISG15, OAS1, STAT1*).
- **Supplementary Figure S2:** Correlation plot of RNA-seq vs qPCR fold-change values for metabolism-related genes (*HK2, GMPR, ENO2, MDH1B, ATP6V1B1*).

**Table S1.** Excel file containing DEG lists from RNA-sequencing analysis across SARS-CoV-2 variants, related to Figures 1 and 3.

**Table S2.** Normalised Enrichment Scores (NES) from fast Gene Set Enrichment Analysis (fGSEA) of oxidative phosphorylation (OXPHOS), cytosolic and mitochondrial metabolic pathways across SARS-CoV-2 variants and time points. NES, adjusted *p*-values (FDR), and leading-edge gene counts are reported for each pathway.

**Table S3.** Excel file containing phospho-kinase array raw and processed data, related to Figure 5.

**Table S4.** Excel file containing raw and normalised intracellular amino acid levels (nmol/10⁶ cells), related to Figure 6.

**Table S5:** Primer sequences and RefSeq accession numbers used for qPCR validation.

## MATERIALS AND METHODS

### Cells and viruses

Human nasal airway epithelial cells (NEC)s, maintained at the air–liquid interface (MucilAir™, pooled from 14 donors, catalog no. EP02MP), were obtained in three batches from Epithelix and cultured in MucilAir™ cell culture medium (Epithelix) at 37 °C with 5% CO₂. SARS-CoV-2 variants were propagated and titrated as previously (Ogunjinmi *et al*., 2024; Zhang *et al*., 2022) in African green monkey kidney (VeroE6) cells expressing human angiotensin-converting enzyme 2 (ACE2) and transmembrane protease serine 2 precursor (TMPRSS2) (VAT), kindly provided by the MRC-University of Glasgow Centre for Virus Research, Glasgow (Rihn *et al*., 2021). VAT cells were maintained in Dulbecco’s Modified Eagle Medium (DMEM) supplemented with 10% fetal calf serum (FCS), 1 mg/mL Geneticin (Gibco), and 0.2 mg/mL Hygromycin B (Invitrogen). Viral stocks were generated and titrated in VAT cells for the following SARS-CoV-2 variants: wild-type/D614G (hCoV-19/England/IC19/2020, B.1.13, EPI_ISL_475572), Alpha (hCoV-19/England/205080610/2020, B.1.1.7, EPI_ISL_723001), Delta (hCoV-19/England/SHEF-10E8F3B/2021, B.1.617.2, EPI_ISL_1731019), Omicron BA.1 (hCoV-19/England/M21021166/2022, BA.1), and BA.5 (hCoV-19/England/NWLP_53/2022, BA.5).

### RNA-sequencing of nasal airway epithelial cells infected with SARS-CoV-2 variants

Before infection, NEC cultures were washed with serum-free medium to remove mucus and debris. Cells were then infected with 200 μL of virus-containing serum-free DMEM (MOI: 0.01) and incubated at 37 °C for 1 hour. Following incubation, the inoculum was removed, and the cells were washed twice. Time points were collected by adding 200 μL of serum-free DMEM, incubating at 37 °C for 10 minutes, and then removing the medium for titration. Cells were harvested at 24 and 72 hours for RNA isolation using the RNeasy Mini Plus kit (Qiagen), and approximately 1 μg of RNA was used for sequencing library preparation.

mRNA libraries were prepared using poly-A enrichment and sequenced by Novogene. The sequencing data (FASTQ raw files) were imported into Partek Flow (version 10.0, build 10.0.23.0531; Partek Inc.) for quality control and processing. Paired-end reads were trimmed based on a Phred quality score threshold of >20 and aligned to the human genome (hg38) using the STAR-2.7.8a aligner. Gene expression was quantified using the Ensembl Transcripts release 104 v2 transcript model, and data were normalised using the counts per million (CPM) method. Differential gene expression analysis was conducted using the DESeq2 package, with genes considered differentially expressed if they met the criteria of FDR ≤ 0.05 and a log2 fold change ≥ |1|. All RNA-seq data have been deposited in the Gene Expression Omnibus (GEO) under accession number GSE271378.

### Pathway enrichment and transcriptomic profiling

Normalised enrichment scores (NES) generated from the fGSEA analysis were visualised using dot plots in R (v4.3) with the *ggplot2* (v3.4.4), *dplyr*, *tidyr*, *stringr*, and *readxl* packages. Each dot represented a metabolic pathway under a specific condition, where the horizontal axis denoted NES magnitude and the vertical axis listed the curated gene sets (cytosolic, mitochondrial, and oxidative phosphorylation). Dot colour indicated the direction of regulation (red, up-regulated; blue, down-regulated), and dot size was proportional to the absolute NES value. Vertical reference lines at NES = 0 were included to indicate neutral enrichment.

To aid visual interpretation, NES magnitudes were categorised as limited (|NES| < 1.0), moderate (1.0 ≤ |NES| < 1.8), or strong (|NES| ≥ 1.8) enrichment. Only pathways meeting the inclusion criteria defined in the differential expression analysis were displayed. While the dot plots provide a comparative overview of pathway-level regulation, they represent aggregate enrichment statistics rather than individual gene contributions. Adjusted *p*-values and false discovery rates (FDR) computed during fGSEA were not visualised in the figure but are reported in Supplementary Table S2. Pathway sizes (number of genes per set) and inter-gene variance were not displayed to maintain visual clarity; therefore, pathways of different gene-set sizes appear equally weighted. Similarly, replicate-level variability and confidence intervals for NES values were not plotted, as the scores reflect ranked enrichment rather than absolute expression magnitude. Relationships between overlapping pathways were not visualised but were accounted for during pathway curation. Together, these dot plots summarise the direction and magnitude of metabolic pathway enrichment across SARS-CoV-2 variants and time points while maintaining a uniform visual scale for comparative assessment.

### Quantification of viral and host gene expression by RT-qPCR

Virus genome quantification was carried out using *E* gene RT-qPCR, following a previously described method (Zhou *et al*., 2022). Viral RNA was extracted from swab supernatant samples using the QIAsymphony DSP Virus/Pathogen Mini Kit on the QIAsymphony instrument (Qiagen). RT-qPCR was performed with the AgPath RT-PCR kit (Life Technologies) on a QuantStudio 7 Flex Real-Time PCR System, using primers specific to the SARS-CoV-2 *E* gene (Corman *et al*., 2020). For absolute quantification of *E* gene RNA copies, a standard curve was generated from serial dilutions of viral RNA with a known copy number. The *E* gene copies per ml of the original virus supernatant were then determined based on this standard curve.

### Validation of RNA-sequencing data by quantitative RT-PCR

To validate differential gene expression observed in RNA-seq, we performed quantitative RT-PCR (qPCR) using independently designed primers as before (Giotis et al., 2019; Ogunjinmi *et al*., 2024; Zhang *et al*., 2022). Genes were selected based on their transcriptional regulation across SARS-CoV-2 variants and their relevance to innate immune signalling (e.g., *CXCL10, IFIT1, IFIT2*, *ISG15*, *OAS1, STAT1*) or metabolic pathways (e.g., *HK2*, *GMPR, MDH1B*, *ENO2, ATP6V1B1*). Total RNA was extracted from nasal epithelial cell cultures using TRIzol™ Reagent (Fisher Scientific) according to the manufacturer’s protocol. qPCR was carried out on a CFX96 Real-Time PCR System (Bio-Rad) using PowerUp™ SYBR Green Master Mix (Fisher Scientific). All reactions were run in triplicate to ensure reproducibility. Primer sequences, designed based on NCBI RefSeq transcripts, are listed in Table S5.

### Phospho-kinase array and signalling analysis

Phosphorylation profiling was carried out using the Proteome Profiler Human Phospho-Kinase Array Kit (ARY003B, R&D Systems), which detects phosphorylation at 43 kinase sites across multiple host signalling pathways. Arrays were performed on lysates collected at 24 hpi from the same nasal epithelial cell cultures used for transcriptomic and amino acid profiling, ensuring matched experimental conditions across assays. Following protein extraction, total protein was quantified using the Pierce BCA Protein Assay Kit (Thermo Scientific), and equal amounts of lysate were incubated with array membranes overnight at 4 °C, according to the manufacturer’s instructions. Chemiluminescent signals were developed using SuperSignal West Femto substrate (Thermo Scientific) and imaged with the Invitrogen iBright FL1500 system. Spot intensities were quantified in ImageJ (v1.53), corrected for background, and normalised to internal reference spots. Fold changes were calculated relative to mock-infected controls. Each condition was assessed in at least two independent experiments. Data visualisation was performed in Python (v3.10) using seaborn and matplotlib. Phosphorylation sites were grouped by pathway (e.g., PI3K-Akt, MAPK, JAK-STAT) based on KEGG and Reactome annotations (Figure 5). Dot plots represent mean fold changes ± SEM, with the mock baseline shown as a red dashed line.

### Quantitative amino acid profiling and network-based visualisation of metabolic disruption

Amino acid analysis was performed using hydrophilic interaction chromatography-tandem mass spectrometry (HILIC-MS/MS), similar as previously described (Jeffares *et al*., 2015; Mulleder *et al*., 2017; Rallis *et al*., 2021). We used four biological replicates per condition, each processed in duplicate (two technical replicates), to ensure robustness and reproducibility of quantification. One million cells were harvested per condition (mock and viral infections). For extraction, cells were pelleted (3000g, 3 min), washed briefly, and resuspended in 200 µL of 80% ethanol preheated to 80 °C. The suspension was vortexed and incubated at 80 °C for 2 minutes, repeated once, then centrifuged at maximum speed for 2 minutes to clear cell debris and stored at 80 °C until further usage. For the exact amino acid quantification, we included cell free ^15^N^13^C labelled amino acids as internal standard (CNLM-6696-PK, Cambridge Isotope Laboratories). Besides, a calibration curve was generated using serial dilutions of standards. The amino acid measurements were performed by using a flowrate of 0.6 ml/min for 12 min. All proteinogenic amino acids amino expect cysteine were measured.

Final amino acid concentrations were expressed as nanomoles (nmol) per 10⁶ cells to normalise for input cell number across conditions (Table S4). To visualise SARS-CoV-2 variant-specific amino acid expression changes, a bipartite node network was constructed using Python (pandas, numpy, networkx, matplotlib). Amino acid log₂ fold changes relative to mock at 24 hpi (Table S4) were used to identify the top five most dysregulated amino acids per variant (ranked by absolute log₂ fold change). These variant–amino acid pairs formed the network edges. Edge thickness reflected the magnitude of the change, while colour indicated direction: red (increased abundance), blue (decreased abundance), and black (no substantial change). Amino acid nodes were sized by total connectivity and coloured according to their dominant direction of change across variants. The layout was manually optimised for clarity. This network allowed integrated visualisation of shared and distinct metabolic disruptions across variants, complementing the quantitative log₂ fold-change data presented in heatmaps.

## Statistical analysis

All data were analysed using GraphPad Prism software (v9) and Python (v3.10). Results are presented as mean ± SEM. Statistical significance was defined as p < 0.05, with levels of significance indicated as follows: p < 0.05 (\**), p < 0.01 (*\*\**), and p < 0.001 (*\*\*\*).

To assess differences in viral replication between SARS-CoV-2 variants at each time point relative to the reference strain IC19, a one-way ANOVA was performed at 0, 24, 48, and 72 hpi. If statistical significance (p < 0.05) was observed, Tukey’s Honest Significant Difference (HSD) test was conducted for post-hoc pairwise comparisons to determine specific differences between IC19 and each variant. Adjusted p-values from Tukey’s HSD test were used to control for multiple comparisons, ensuring robust identification of significant replication differences. Variants exhibiting p-values below the 0.05 threshold were considered to have significant replication advantages or disadvantages relative to IC19. Results were annotated on viral growth curves to highlight meaningful differences across the infection timeline.

Phospho-array data were log-transformed data and analysed using Welch’s t-tests, followed by Benjamini-Hochberg correction for multiple comparisons (Table S4). Amino acid concentrations (nmol per 10⁶ cells) were measured in four biological replicates per condition, each processed in duplicate. Technical replicates were averaged per biological replicate before statistical analysis. For each amino acid, mean values were used to calculate the fold change relative to mock and the corresponding log₂ fold change (log₂FC = log₂[variant/mock]). Statistical testing was performed using Welch’s two-sample t-tests (unequal variances) applied to each amino acid for: (i) comparisons of each SARS-CoV-2 variant against mock, and (ii) all pairwise comparisons between conditions (mock, IC19, Delta, BA.1, BA.5). Multiple testing within each comparison was corrected using the Benjamini–Hochberg false discovery rate (FDR) procedure (threshold q < 0.05). Table S4 reports, for each amino acid and comparison, the raw p value, the FDR-adjusted q value, a Yes/No column indicating significance at q < 0.05, and p-value significance symbols (* p < 0.05; ** p < 0.01; *** p < 0.001; **** p < 0.0001; ns for non-significant). Log₂FC values are provided as effect sizes.

To assess whether infection altered amino acid composition independently of total abundance, amino acid levels were normalised to the total amino acid content within each condition. The resulting relative proportions were compared across variants using the Friedman test with Dunn’s multiple-comparison correction (GraphPad Prism v9). Total amino acid abundance per condition was calculated by summing per–amino acid means, with percent change expressed relative to Mock.

## Data visualisation

To illustrate shifts in amino acid levels across SARS-CoV-2 variants, heatmaps and summary tables were used to facilitate interpretation and guide RNA-seq pathway analysis. For phosphoproteomic analyses, heatmaps were generated using Python’s matplotlib and seaborn libraries. Heatmap of amino acid fold changes were generated using the pheatmap package in R (v4.3), clustering both rows and columns to reveal variant-specific metabolite patterns. Venn diagrams were created with InteractiVenn (Heberle *et al*., 2015) to analyse shared and unique features across different conditions.

