## Supplementary Figures 1 & 2 for "Evolutionary rewiring of host metabolism and interferon signalling by SARS-CoV-2 variants"

**Supplementary material**

**Supplementary Figure 1:** **Correlation between RNA-seq and qPCR expression data for selected immune-signalling-related genes.** Scatter plot comparing log₂ fold-change values from RNA-sequencing and qPCR for selected immune signalling-related genes (CXCL10. IFIT1, IFIT2, ISG15, OAS1, STAT1). Scatter plot comparing log₂ fold-change values obtained from RNA-sequencing and qPCR for **39 values** analyzed under identical experimental conditions. Each point represents a gene. Pearson correlation analysis showed a **very strong positive correlation** (**r = 0.9656, R² = 0.9325**) with a **95% confidence interval of 0.9350 to 0.9820**. The correlation was statistically highly significant (**p < 0.0001**, two-tailed; ****), confirming the robustness and accuracy of RNA-seq in capturing gene expression changes relative to qPCR.

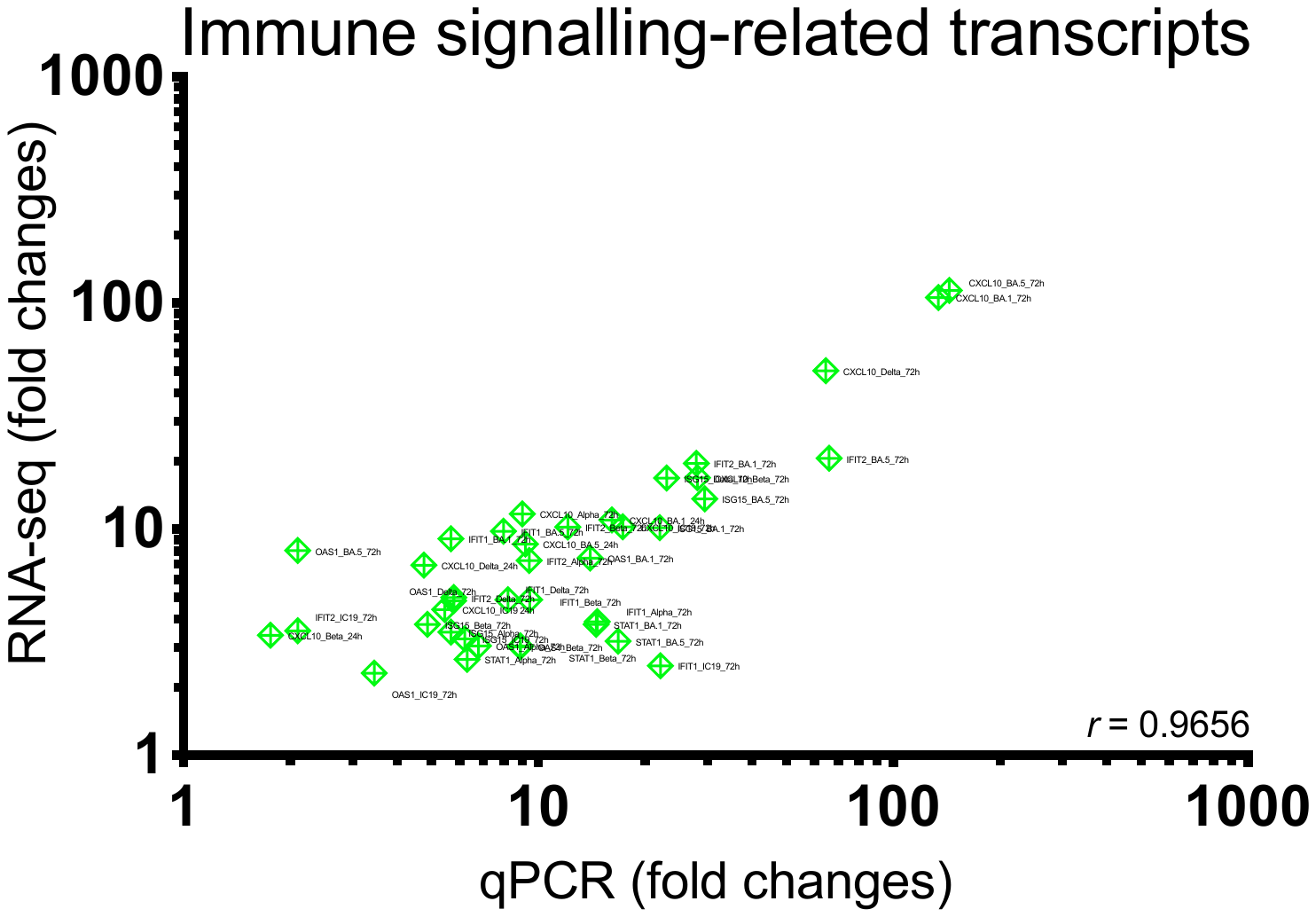

**Supplementary Figure 2: Validation of RNA-seq data by qPCR for selected genes.**
Scatter plot showing the correlation between log₂ fold-change values obtained from RNA-seq and qPCR analysis across selected metabolism-related genes (HK2, GMPR, ENO2, MDH1B, ATP6V1B1). Correlation analysis between log₂ fold-change values obtained from RNA-sequencing and qPCR for 37 values. Each point represents one gene. Pearson correlation analysis revealed a strong positive correlation (**r = 0.9310, R² = 0.8668**) with a **95% confidence interval** of 0.8692 to 0.9642, indicating high consistency between the two methods. The correlation was statistically significant (**p < 0.0001**, two-tailed; ****), confirming the robustness and accuracy of RNA-seq in capturing gene expression changes relative to qPCR.

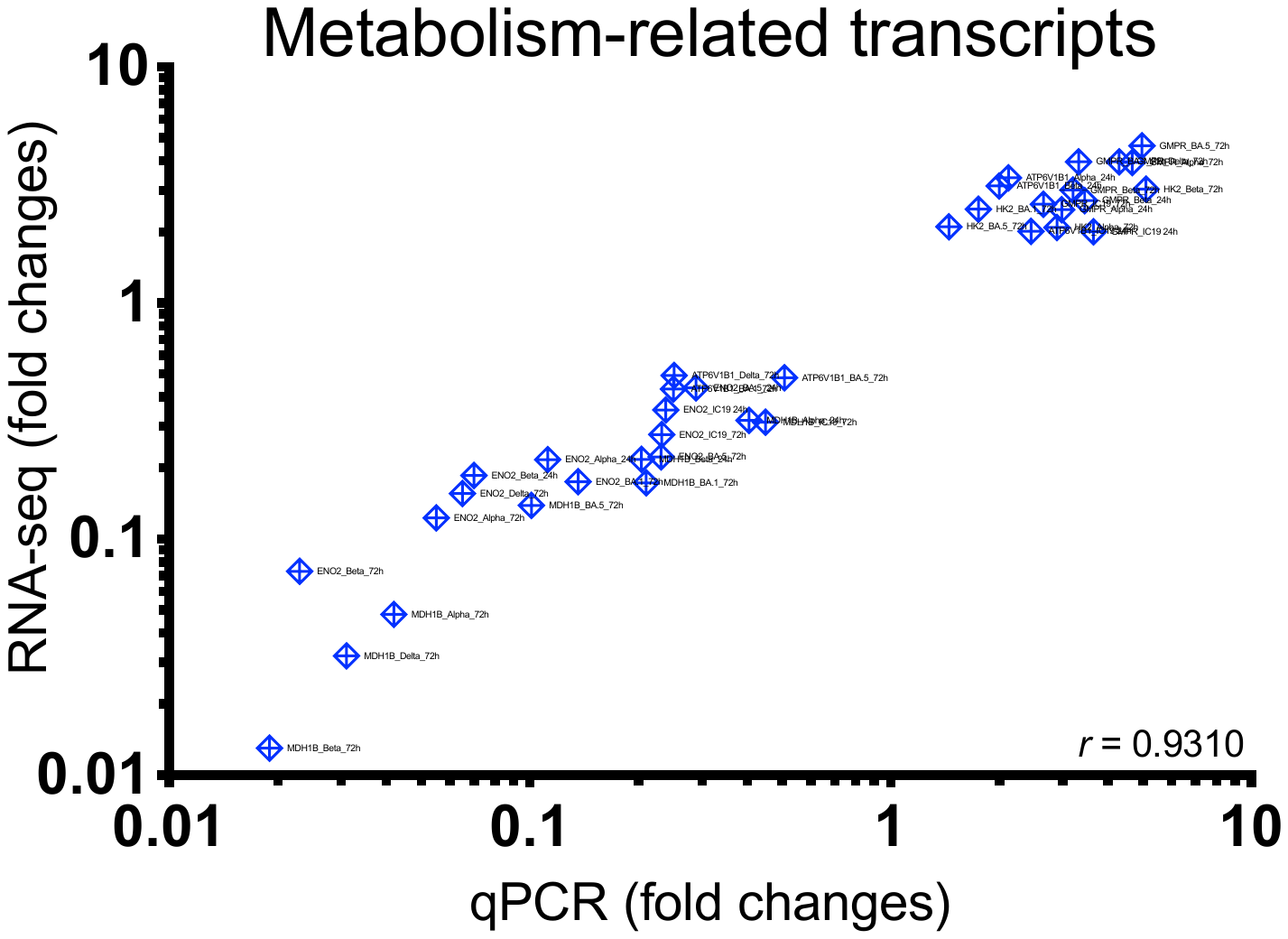
