## Supplementary Table 5 for "Evolutionary rewiring of host metabolism and interferon signalling by SARS-CoV-2 variants"

| **Gene** | **RefSeq Accession Number** | **Forward Primer** | **Reverse primer** |
| --- | --- | --- | --- |
| **GAPDH** \| \| | \| NM_002046 \| | AATCCCATCACCATCTTCCA | TGGACTCCACGACGTACTCA |
| **CXCL10** | \| NM_001565 \| | TCTCTCTAGAACTGTACGCTGT | ACATGATCTCAACACGTGGACA |
| **CXCL11** \| \| | \| NM_005409 \| | AGCAGTGAAAGTGGCAGAT | TTGGGATTTAGGCATCGT |
| **IFIT1** \| \| | \| NM_001548 \| | GCCTTGCTGAAGTGTGGAGGAA | ATCCAGGCGATAGGCAGAGATC |
| **IFIT2** \| \| | NM_001547 \| | CAGCTGAGAATTGCACTGCAA | CGTAGGCTGCTCTCCAAGGA |
| **ISG15** \| | \| NM_005101 \| | GGACAAATGCGACGAACCTC | GAGGTTCGTCGCATTTGTCC |
| **IRF7** \| \| | \| NM_001572 \| | AGCTGCACGTTCCTATACGG | CCGTATAGGAACGTGCAGCT |
| **OAS1** \| \| | \| NM_016816 \| | AGGAAAGGTGCTTCCGAGGTAG | GGACTGAGGAAGACAACCAGGT |
| **MX2** \| | NM_002463 \| | AGCCACCACCAGGAAACA | TTCTGCTCGTACTGGCTGTACAG |
| **IFI44** \| \| | \| NM_006417 \| | CGGTTGCACGAAAAGATCCTG | GTCAAGCAAAACTCCATTACGGA |
| **RSAD2** \| \| | \| NM_080657 \| | TGGGTGCTTACACCTGCTG | TGAAGTGATAGTTGACGCTGGT |
| **IFITM3** \| \| | NM_021034 \| | CTGGGCTTCATAGCATTCGCCT | AGATGTTCAGGCACTTGGCGGT |
| **STAT1** \| \| | \| NM_007315 \| | ACACGAGACCAATGGTGTGG | AACTTGCTGCAGACTCTCCG |
| **HK2** \| \| | \| NM_000189 \| | GAGTTTGACCTGGATGTGGTTGC | CCTCCATGTAGCAGGCATTGCT |
| **GMPR** \| \| | \| NM_006877 \| | TGAACAAGCACGCAGGAGGA | AACACGGTGTTGTGCTGCTG |
| **ENO2** \| \| | \| NM_001975 \| | CTGTATCGCCACATTGCTCAGC | AGCTTGTTGCCAGCATGAGAGC |
| **MDH1B** \| \| | \| NM_206892 \| | CAACCCCTTGCAGGTCTGGA | GTGCAGCCAAAATGCCTCCA |
| **ATP6V1B1** \| | \| NM_001692 | CGGGTTTTCAATGGCTCCGG | GCGGCAATCTCATTGTGGGG |
